# Host plant odour and sex pheromone are integral to mate finding in codling moth

**DOI:** 10.1101/2023.10.02.560528

**Authors:** Anna Erdei, Michele Preti, Marie Bengtsson, Peter Witzgall

## Abstract

Behaviour-modifying chemicals mediate sexual communication and host choice in insect herbivores. Sex pheromones are believed to attract insects by themselves, even though they are released into an atmosphere of plant odorants. We show for the first time that, in codling moth, feeding on apple and pear, that female pheromone is efficient for male attraction only in the presence of host plant odour. In non-host vegetation, male attraction to sex pheromone was very strongly reduced. The role of host odour in sex attraction was then substantiated by blending synthetic pheromone, codlemone, and the kairomone pear ester, a strong host plant attractant. An admixture of pear ester entirely rescued pheromone attraction in non-host vegetation. This field behavioural assay substantiates that host plant olfactory cues are integral part of sexual communication and mate recognition, which provides a mechanism for how shifts to new host plants produce new mate recognition signals.

## Introduction

The extraordinary diversity of plant-feeding insects depends on specific host plant associations and in consequence on host plant recognition. Insects, and particularly nocturnal insects, use first of all the sense of smell to find their plant hosts. Plant odorants mediate behavioural responses directed to the plant, like feeding and oviposition, but they also contribute to specific mate communication, in concert with sex pheromones, since matings often occur on the host plant (Ehrlich & Raven 1964; Smadja & Butlin 2009; Janz 2011; Borrero-Echeverry et al. 2018).

Taxonomically related insects are often found on taxonomically related plants and this indicates that ecological divergence may even occur in sympatry. Mating preferences coupled to habitat or host plant choice are indeed thought to play a role in speciation, a combination of sexual and natural selection is probably sufficient to initiate and complete speciation (Blows et al. 2002; Bolnick & Fitzpatrick 2007; Maan & Seehausen 2011; Butlin et al. 2012; Scordato et al. 2014; Rosenthal 2017).

The concept that habitat and mate choice are linked has since long been established, while the underlying sensory mechanisms are not yet fully resolved. In codling moth *Cydia pomonella*, molecular and physiological evidence strongly supports the idea that configural blends of sex pheromone and host plant volatiles are behaviourally particularly powerful.

The larvae of codling moth feed on apple, but also on walnut, pear or other rosacean fruits and codling moth is economically very important in all fruit growing areas. The female sex pheromone codlemone (Arn et al. 1985) and pear ester, a kairomone or host plant attractant (Light et al. 2001; Light & Knight 2005), are widely used for environmentally safe control (Witzgall et al. 2008; Knight et al. 2018).

The kairomone pear ester is perceived via an olfactory receptor that belongs to the clade of pheromone receptors (Bengtsson et al. 2014; Wan et al. 2019) and in male moths, a blend of pheromone codlemone and pear ester elicits a conspicuously synergistic response in the antennal lobe, the olfactory centre in the insect brain (Trona 2010, 2013).

We here show an elementary field attraction assay that complements the molecular and physiological evidence and that conclusively demonstrates the deep link between the perception of social and environmental olfactory signals. Female sex pheromone elicits the male attraction response only in the presence of host plant odour.

In nature, pheromone is released into a background of plant odorants. Our field behavioural assay demonstrates that the response to individual semiochemicals cannot be singled out and that sex signals and habitat cues are integral components of mate-finding. It follows that their perception is under combined sexual and ecological selection.

## Methods and materials

Codlemone, (*E,E*)-8,10-dodecadienol (E8,E10-12OH), a gift from K. Ogawa (Shin-Etsu Chemical Co., Chiyoda, Japan), was >99.6% pure. Pear ester, (*E,Z*)-2,4-decadienoate (Sigma Aldrich), was ≥95% pure (chemical and isomeric). Compounds were diluted in hexane (Sigma Aldrich, redistilled) and formulated on red rubber septa (Merck ABS, Dietikon, Switzerland), which were used as field lures during up to 2 wk. Lures contained codlemone, pear ester, or a blend of both compounds, they were mounted in tetra traps (Arn et al. 1979).

Traps were placed in orchards, near Kivik (Scania, Sweden), where rows of apple trees (*Malus domestica*) are interspersed with rows of other trees that serve as windbreaks, such as birch (*Betula pubescens*) or whitebeam (*Sorbus aria*). These windbreaks were directly adjacent to rows of apple, on the left and right side. Windbreak trees do not support development of codling moth larvae. In some orchards, blocks of several rows of apple trees are adjacent to blocks of other fruit trees, such as pear (*Pyrus communis*), a codling moth host plant, or plum (*Prunus domestica*).

Traps were placed in apple trees, in rows that were either adjacent to wind breaks or adjacent to blocks of other fruit trees, and the same number of traps was placed in windbreaks or in fruit trees directly adjacent to apple. Distance of apple tree stems was 4 to 5 m from tree stems in adjacent rows of fruit trees or non-host trees serving as windbreaks.

Further test were done near Faenza (Ravenna, Italy) in pear orchards with an adjacent peach orchard or an adjacent vineyard. Traps were placed in pear trees and in adjacent peach trees or in vines, respectively.

Traps were hung at 1.5 to 2 m from the ground to green branches, and witnessed every 2 to 3 days. For statistical analysis, captures in apple and the corresponding captures in other fruit trees or windbreaks were compared using a two-tailed Student’s t-test.

## Results

Attraction of codling moth males to traps baited with the female sex pheromone codlemone declined when traps were placed in trees other than apple, even though these trees were directly adjacent to apple (Figure 1 A). Experiments were conducted in apple orchards, in apple tree rows next to rows of other fruit trees or windbreak rows, with tree stems 4 to 5 m apart.

**Figure 1.**
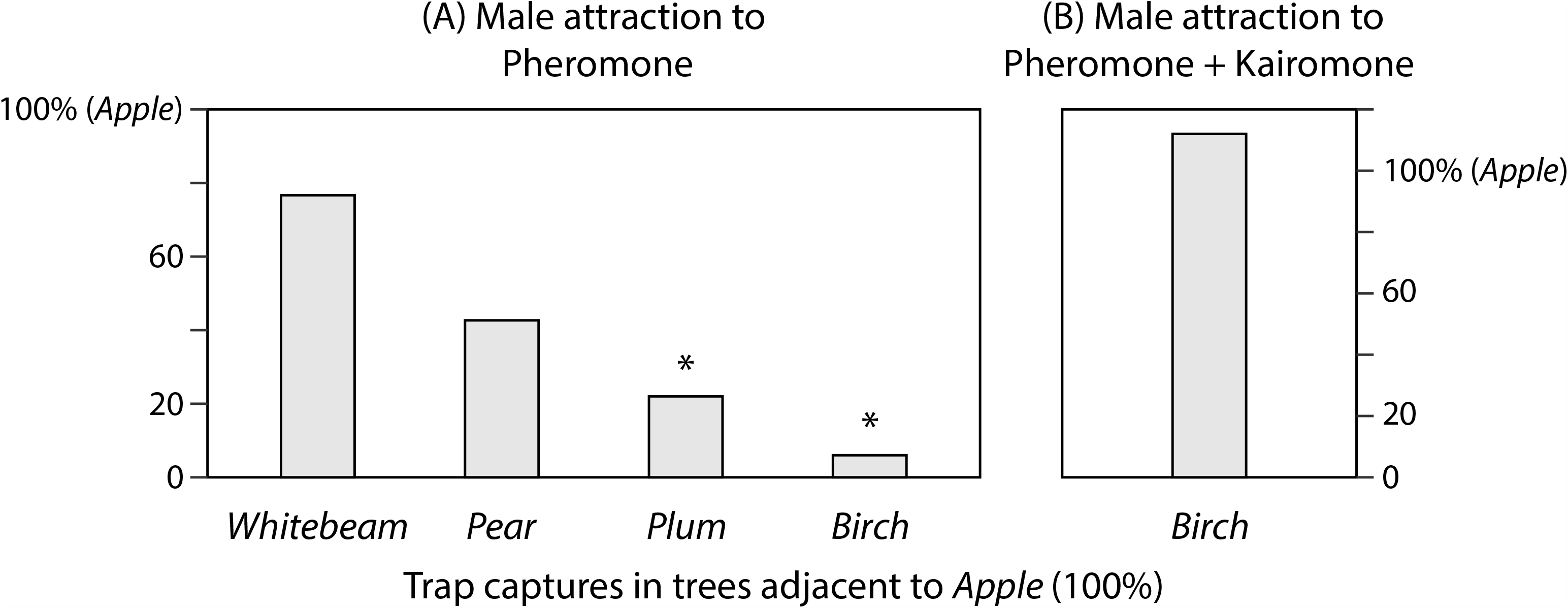
Codling moth *Cydia pomonella* male captures in traps baited with codling moth sex pheromone (codlemone), and a blend of pheromone and a kairomone or host plant attractant (pear ester). (A) Captures of male moths with 100 µg of sex pheromone codlemone in four apple orchards, in trap pairs placed in apple (100%) and in adjacent windbreak rows or other fruit trees, whitebeam *Sorbus aria* (n=9), pear *Pyrus communis* (n=9), cherry *Prunus avium* (n=11), or birch *Betula pubescens* (n=6). Captures in cherry and birch were significantly lower than in corresponding traps in apple (t=2.9163, p=0.008537 and t=5.98661, p=0.000134, respectively, two-tailed t-test). (B) Male moth captures with a blend of 100 µg sex pheromone codlemone and 100 µg of the kairomone pear ester, in apple and adjacent birch trees were not significantly different (n=50, t=0.69816, p=0.486733, two-tailed t-test).

In apple orchards in Sweden, differences between captures in apple and the non-host plants cherry (*Prunus avium*, Rosaceae) and birch (*Betula pubescens*, Betulaceae) were significant. Captures in pear (*Pyrus communis*) and whitebeam (*Sorbus aria*, both Rosaceae) were not significantly lower than in adjacent apple trees. Codling moth females also oviposit on pear, while there are no records from whitebeam (Figure 1 A).

In pear orchards in Italy, differences between male captures in pear and adjacent peach trees were not significant, peach is an occasional host of codling moth. The ratio of the number of males captured in peach vs. pear was 1.17 (n=10 traps, n=76 males; t=-0.33007, p=0.74516). In comparison, captures with codlemone significantly declined in grapevine, which is not a suitable host. The ratio of the number of males captured in grape vs. pear was 0.09 (n=10 traps, n=130 males; t=2.98039, p=0.00802; two-tailed t-test).

We then aimed to verify the remarkable discrepancy in captures in apple and adjacent birch windbreak rows with a dose-response test. At 1, 10 and 100 µg codlemone, captures were significantly lower in traps placed in birch, compared to apple (Table 1).

**Table 1.** Codling moth *Cydia pomonella* males trapped with 1, 10 and 100 µg of codling moth sex pheromone (codlemone), and with a kairomone or host plant attractant (pear ester), in apple and and an adjacent windbreak row of birch trees, within an apple orchard. Captures with codlemone in apple and birch were significantly different (p<0.05, n=6; two-tailed t-test). Pear ester alone is a weak attractant, captures in apple and birch were not significantly different.

| Trap placement | Codlemone | 1 | 10 | 100 µg |
| --- | --- | --- | --- | --- |
| Apple |  | 0.71 a | 1.51 a | 4.44 a |
| Birch |  | 0 b | 0.04 b | 0.28 b |
|  | Pear Ester | 1 | 10 | 100 µg |
| Apple |  | 0 | 0.07 | 0.14 a |
| Birch |  | 0 | 0 | 0.03 a |

Remarkably, the decline in trap captures in birch trees was entirely restored by blending sex pheromone with the kairomone or host plant attractant pear ester (Figure 1 B). Pear ester by itself is a weak attractant. A dose-response test with pear ester alone confirmed that few males were attracted to pear ester alone, in apple and birch (Table 1), while it had a significant effect on male attraction in birch, when blended with female sex pheromone (Figure 1 B).

## Discussion

### Sex pheromone and host odour mediate mate finding in codling moth

Our field study affords compelling evidence that chemosensory attraction of codling moth males to females is mediated by a blend of sex pheromone and host plant odour, and that both are integral components of mate recognition and mate finding (Borrero-Echeverry et al. 2018). It follows that divergence onto new host plants generates new premating communication channels that may give rise to reproductive isolation.

This corroborates the concept that mating preferences coupled to habitat choice lead to an interaction between sexual and ecological selection, which plays a powerful role in ecological divergence and speciation (Bolnick & Fitzpatrick 2007; Ritchie 2007; Schluter 2009; Maan & Seehausen 2011; Servedio et al. 2011; Butlin et al. 2012; Debelle et al. 2014; Rosenthal 2017; Scordato et al. 2014; Blows 2022)

Codling moth responses to sex and host plant signal cannot be dissociated, which makes it difficult or even obsolete to distinguish sexual from natural selection. Furthermore, powerful synergistic and antagonistic interactions between social and plant olfactory signals, in combination with a patchy distribution of host trees in natural habitats, is expected to generate assortative mating and to restrict gene flow even among species found in the same geographic region. The distinction of sympatry vs. allopatry does accordingly not apply to specialist herbivores where host choice is combined with mate choice (Mallet et al. 2009).

Field observations invariably show that codling moth females are not evenly distributed in apple orchards, but tend to aggregate around prominent trees with strong fruitsetting, probably due to a combination of olfactory and visual cues. The males are on their wings and seen to fly around the crowns of these trees well before the onset of female pheromone emission (Witzgall et al. 1999; Bengtsson et al. 2001). This illustrates that the male response to host plant volatiles is also under sexual selection. That male moths fly about their host plants, before the onset of the diel mating period, has been observed in many other tortricid moths (Bradley et al. 1979).

As soon as codling moth females start to release sex pheromone, closeby males immediately respond, which is why pheromone-releasing females are very rarely seen in the field (Witzgall et al. 1999). In comparison, in the laboratory, isolated females maintain the typical calling posture, lifted abdomen and extended ovipositor, during several hours (Bäckman et al. 1997).

### Blends of sex pheromone and host plant odorants encode species-specificity

The combined response to sex pheromone and host plant volatiles is also under natural selection, since the host plant plays a prominent heterospecific role in reproductive isolation. The few *Cydia* species that use the same sex pheromone are all found on taxonomically widely distant hosts (Witzgall et al. 1996). For example, pear moth *C. pyrivora*, feeding on wild and cultivated pear (Rosaceae) and pea moth *C. nigricana*, feeding on peas and vetches (Fabaceae), both use the same sex pheromone, (*E,E*)-8,10-dodecadienyl acetate (codlemone acetate) (Witzgall et al. 1993; Makranczy et al. 1998).

On the other hand, pear moth and codling moth are sibling species that both co-occur on pear. Codling moth attraction to its sex pheromone codlemone is strongly reduced in the presence of pear moth pheromone, codlemone acetate, and vice versa (Witzgall et al. 1996; Witzgall et al. 2010). Codling moth males express an olfactory receptor (OR) in antennal, olfactory sensory neurons (OSNs) that is specifically tuned to codlemone acetate (Bäckman et al. 2000, Walker et al. 2016, Cattaneo et al. 2017) and that evidently serves perception of heterospecific pheromone.

Codling moth feeds on apple and pear in Northern Europe and also on walnut in Southern Europe or overseas, and there are no apparent pheromone dialects among these host plant populations (Dumenil et al. 2014). In comparison, host plant races of several other moths use different pheromone blends, such as European corn borer *Ostrinia nubilalis* and larch budworm, *Zeiraphera diniana* (Priesner & Baltensweiler 1987; Pelozuelo et al. 2004; Malausa et al. 2005). However, assortative mating among these host races does not rely on different pheromone signatures alone (Pelozuelo et al. 2007). A contributing role of the host plant has been shown in *Z. diniana*, where females attract males from within the same or adjacent trees (Emelianov et al. 2001), which aligns with our obserrvations in codling moth. In cotton leafworm *Spodoptera littoralis*, attraction to heterospecific pheromone, of the sibling species *S. litura*, is greatly reduced in the presence of the host plant (Borrero et al. 2018). Likewise, mate preference phenotypes covary among host plant genotypes in in hemipteran treehoppers (Rebar & Rodriguez 2015).

### Essential roles of plant volatiles in pheromone communication and reproductive behaviour

Plants and plant odorants strongly impact the reproductive behaviour of lepidopteran insects. The physiological and behavioural effects span from sexual maturation, feeding, enhanced sex attraction and courtship to oviposition. Especially the effect of plant odorants on the male response to female sex pheromone has received attention, and both synergistic and antagonistic effects have been found (Landolt & Phillips 1997; Reddy & Guerrero 2004; Gadenne et al. 2016; Borrero-Echeverry et al. 2018; Conchou et al. 2019)

Seemingly conflicting effects are sometimes due to changes in the reproductive state of the insect, or the condition of the plant. For example, in cotton leafworm, males are attracted to larval host plants as rendez-vous sites prior to mating, and to floral volatiles for feeding post mating. Both males and females avoid host plants damaged by feeding larvae, in response to defense-induced plant volatiles (Zakir et al. 2013; Hatano et al. 2015; Kroman et al. 2015).

A most serious shortcoming is that the key plant compounds that mediate specific host attraction are only known for a handful of species, also because live plants are not easily manageable in controlled laboratory experiments. Many plant volatiles have been found to mediate some level of attractation in many insect species, but these include ubiquitous compounds, produced by a wide range of plants, that cannot account for specific host plant attraction. Plants also release very many different volatiles, their abundance does not correlate with their behavioural activity in insects, and we lack straightforward techniques for reliable identification (Gonzalez et al. 2020).

Our results show that the very essential role of the host plant in pheromone communication in insect herbivores has not been fully realized, where the incomplete knowledge of the key bioactive plant compounds has been a contributing factor. Our radically simple approach to place pheromone-baited traps at close distance in non-host trees bypasses this shortcoming, and impressively demonstrates, for the first time, the paramount role of the host plant in pheromone communication.

Hundreds of lepidoperan pheromones have been identified (El-Sayed 2023), through chemical analysis followed by a field trapping bioassay. These traps have been routinely placed in the right habitat, on the respective host plants, but attraction has been attributed to the pheromone lures only.

### Antagonistic effect of non-host plant volatiles

Trap captures in different trees seemingly reflect taxonomic relatedness and chemical similarity with apple. Captures were highest in apple and gradually declined among three Rosacean species to birch.

There is some evidence for the occurence of host races in codling moth, and a differential response of these host races to their respective host plants. Apple is the main host worldwide. In Sweden, codling moth feeds also on pear, peach is an occasional host, but codling moth has not been reported from plum (Bovey 1966; Cisneros & Barnes 1974; Phillips & Barnes 1975, Witzgall et al. 2005)

In birch, grapevine or plum, it remains unclear whether the lack of attraction is due to absence of attractants, or rather an antagonistic interference of non-host odorants and codling moth pheromone, which almost annihilates male moth attraction, for example in birch. Hedgerows and windbreaks contribute to codling moth management in orchards, and reduce teh number of codling moth diapausing larvae in their vicinity (Ricci et al. 2011), which points towards an antagonistic effect of non-host plant volatiles. Antagonistic pheromone and non-host plant interactions have been studied, particularly in bark beetles (Byers et al. 1998; Zhang & Schlyter 2003, 2004) and also in moths (McNair et al. 2000; Wang et al. 2016), with respect to insect management.

### Molecular and physiological mechanism of codlemone-pear ester interaction

The neurophysiology and behavioural ecology of pheromone-plant volatile interactions has been thoroughly investigated in moths (Christensen & Hildebrand 2002; Namiki et al. 2008; Deisig et al. 2012; Pregitzer et al. 2013; Chaffiol et al. 2014). However, the knowledge of the key plant volatiles that are perceived via dedicated olfactory channels, to elicit behavioural responses, is a main requirement for investigations of the cognitive architecture of plant-pheromone interactions.

Semiochemicals that mediate plant-insect interactions have been termed “kairomones”, in analogy to pheromones. Such a kairomone has been discovered in codling moth. Pear ester strongly attracts adult males and females, and also larvae (Knight & Light 2001; Light et al. 2001; Light & Knight 2005). Its attractant power enables practical use, for example to monitor the seasonal abundance of codling moth as well as to enhance population control by mating disruption, in blends with codlemone (Knight et al. 2018). That pear ester spectacularly restores pheromone captures of codling moth males in birch underlines that this compund mediates host plant attraction.

Intriguingly, pear ester is perceived via CpomOR3, an odorant receptor (OR) that belongs to the conserved clade of lepidopteran pheromone receptors. CpomOR3 was found in a codling moth antennal transcriptome and its main ligand, according to a functional assay in *Drosophila* olfactory sensory neurons, is pear ester (Bengtsson et al. 2014; Walker et al. 2016; Gonzalez et al. 2017). Male-biased pheromone receptors respond mainly to sex pheromones, while general ORs are tuned to environmental odors including plant volatiles (Fleischer et al. 2018; Ha & Smith 2022; Yang et al. 2022).

A recent codling moth genome assembly reveals a duplication of CpomOR3. A functional assay shows that both copies are tuned to pear ester, and that they also respond to the sex pheromone codlemone (Wan et al. 2019). This adds further evidence to the strong synergy between pheromone, codlemone and kairomone, pear ester and that the odorant receptor CpomOR3 encodes both host plant location and mate finding. Traits that combine ecological and sexual selection are particularly powerful during phylogenetic divergence (Servedio et al. 2011; Merrill et al. 2012; Safran et al. 2013).

In addition to CpomOR3, the synergism between codlemone and pear ester also involves the yet unidentified main pheromone receptor for codlemone, which is expressed in sensilla trichodea, that connect to the macroglomerular complex (MGC) of the antennal lobe (AL) (Bäckman et al. 2000, Trona et al. 2010, 2013). Glomeruli in the AL are the site of convergence for olfactory input from olfactory sensory neurons expressing the same olfactory receptors. Olfactory neurons extend from sensilla on the antenna to glomeruli in the AL. In male moths, the MGC is the area dedicated to processing of pheromone input (Hansson & Anton 2000).

Intracellular recordings from OSNs projecting to the AL, as well as functional imaging of the AL show a very strong synergistic interaction in the MGC, upon stimulation with blends of codlemone and pear ester. Codlemone OSNs project to the MGC and pear ester OSNs to an adjacent glomerulus, which likely facilitates this extraordinary synergistic effect (Trona et al. 2010, 2013). Phylogenetically closely related ORs, with high sequence similarity, often express in OSNs that project to neighboring glomeruli in the AL (Ramdya & Benton 2010).

Taken together, pheromone and kairomone, codlemone and pear ester, are both perceived by pheromone receptors and the architecture of the codling moth olfactory centre and the physiological response provides a mechanism for the intimate interaction of these two compounds, and reflect the outcome of our field experiment.

## Conclusion

The combined molecular, physiological, and behavioural evidence of host plant and sex pheromone communication in codling moth substantiates a deep interaction between mate and host finding. This interaction largely builds on two chemicals, codlemone and pear ester, that yield a strong behavioural synergism and encode host preference and mate recognition.

An exciting aspect for future research is that the cognitive olfactory architecture of pheromone and kairomone communication in codling moth, including phylogenetically related ORs and interlaced signal coding in the insect olfactory centre, is experimentally tractable. It thus becomes possible to investigate the consequences of adaptation to new hosts or sex pheromone signals for assortative mating and reproductive isolation.

A main remaining question is whether the key compounds that encode host recognition are produced by the plant itself or associated microorganisms (Witzgall et al. 2012; Ljunggren et al. 2019). The combined evidence in codling moth and a future comparison of codling moth host races and closely related species is bound to facilitate the identification of these yet unknown host semiochemicals. Tying the key compounds to chemosensory receptors and pathways that encode host preference and mate recognition in concert will help to conceive a plausible scenario of how host shifts give rise to new communication channels.

## Authors’ contributions

A.E., M.P., M.B. and P.W. prepared and carried out the field experiments. M.B. and P.W. conceived the study, P.W. wrote a draft version of the manuscript, all authors contributed to a revision of the manuscript.

## Data accessibility

The data of this study are shown in the manuscript. Raw data are available from the Mendeley Data repository https://doi.org… (Erdei et al. 2023).

## Acknowledgements

This study was supported by the Swedish Research Council Formas (grant 2021-00834), and the Faculty of Landscape Architecture, Horticulture, and Crop Production Science (SLU, Alnarp).

## References

Arn, H., Rauscher, S. & Schmid, A. (1979).Sex attractant formulations and traps for the grape moth Eupoecilia ambiguella Hb. Mitt. Schweizer Entomol. Ges., 52, 49–55.

Arn, H., Guerin, P.M., Buser, H.R., Rauscher, S. & Mani, E. (1985).Sex pheromone blend of the codling moth, Cydia pomonella: evidence for a behavioral role of dodecan-1-ol. Experientia, 41, 1482–1484. (doi:10.1007/BF01950048)

Bäckman, A.-C., Bengtsson, M. & Witzgall, P. (1997).Pheromone release by individual females of codling moth, Cydia pomonella L. (Lepidoptera: Tortricidae). J. Chem. Ecol., 23, 807–815. (doi:10.1023/B:JOEC.0000006412.16914.09)

Bäckman, A.-C., Anderson, P., Bengtsson, M., Löfqvist, J., Unelius, C.R. & Witzgall, P. (2000).Antennal response of codling moth males, Cydia pomonella (L.) (Lepidoptera: Tortricidae), to the geometric isomers of codlemone and codlemone acetate. J. Comp. Physiol. A, 186, 513–519. (doi:10.1007/s003590000)

Bengtsson, M., Bäckman, A.-C., Liblikas, I., Ramirez, M.I., Borg-Karlson, A.-K., Ansebo, L. et al. (2001).Plant odor analysis of apple: antennal response of codling moth females to apple volatiles during phenological development. J. Agric. Food Chem., 49, 3736–3741. (doi:10.1021/jf0100548)

Bengtsson, J.M., Gonzalez, F., Cattaneo, A.M., Montagne, N., Walker, W.B., Bengtsson, M. et al. (2014).A predicted sex pheromone receptor of codling moth Cydia pomonella detects the plant volatile pear ester. Front. Ecol. Evol., 2, 33. (doi:10.3389/fevo.2014.00033)

Blows, M.W. (2002).Interaction between natural and sexual selection during the evolution of mate recognition. Proc. R. Soc. B, 269, 1113–1118. (doi:10.1098/rspb.2002.2002)

Bolnick, D.I. & Fitzpatrick, B.M. (2007).Sympatric speciation: models and empirical evidence. Annu. Rev. Ecol. Evol. Syst., 38, 459–487. (doi:10.1146/annurev.ecolsys.38.091206.095804)

Borrero-Echeverry, F., Bengtsson, M., Nakamuta, K. & Witzgall, P. (2018).Plant odour and sex pheromone are integral elements of specific mate recognition in an insect herbivore. Evolution, 72, 2225–2233. (doi:10.1111/evo.13571)

Bovey, P. (1966).Super-famille des tortricoidea. Pp 456-893 in Balachowsky AS (ed) Entomologie Appliquée à l’Agriculture, Vol. II. Paris: Masson.

Bradley, J.D., Tremewan, W.G. & Smith, A. (1979).British tortricoid moths. Tortricidae: Olethreutinae. The Ray Society, London, pp. 336.

Butlin, R., Debelle, A., Kerth, C., Snook, R.R., Beukeboom, L.W., Cajas, R.F.C. et al. (2012) What do we need to know about speciation? Tr. Ecol. Evol., 27, 27–39. (doi:10.1016/j.tree.2011.09.002.)

Byers, J.A., Zhang, Q.H., Schlyter, F. & Birgersson, G. (1998).Volatiles from nonhost birch trees inhibit pheromone response in spruce bark beetles. Sci. Nat., 85, 557–561. (doi:10.1007/s001140050551)

Cattaneo, A.M., Gonzalez, F., Bengtsson, J.M., Corey, E.A., Jacquin-Joly, E., Montagné, N. et al. (2017).Candidate pheromone receptors from the insect pest Cydia pomonella respond to pheromone and kairomone components. Sci. Rep., 7:41105. (doi:10.1038/srep41105)

Chaffiol, A., Dupuy, F., Barrozo, R.B., Kropf, J., Renou, M., Rospars, J.P. et al. (2014).Pheromone modulates plant odor responses in the antennal lobe of a moth. Chem. Senses, 39, 451–463. (doi:10.1093/chemse/bju017)

Christensen, T.A. & Hildebrand, J.G. (2002).Pheromonal and host-odor processing in the insect antennal lobe: how different? Curr. Op. Neurobiol., 12, 393–399. (doi: 10.1016/S0959-4388(02)00336-7)

Cisneros, F.H. & Barnes, M.M. (1974).Contribution to the biological and ecological characterization of apple and walnut host races of codling moth, Laspeyresia pomonella (L.): moth longevity and oviposition capacity. Environm. Entomol., 3, 402–406. (doi:10.1093/ee/3.3.402)

Conchou, L., Lucas, P., Meslin, C., Proffit, M., Staudt, M. & Renou, M. (2019).Insect odorscapes: from plant volatiles to natural olfactory scenes. Front. Physiol., 10, 972. (doi:10.3389/fphys.2019.00972)

Debelle, A., Ritchie, M.G. & Snook, R.R. (2014).Evolution of divergent female mating preference in response to experimental sexual selection. Evolution, 68, 2524–2533. (doi:10.1111/evo.12473)

Deisig, N., Kropf, J., Vitecek, S., Pevergne, D., Rouyar, A., Sandoz, J.C. et al. (2012).Differential interactions of sex pheromone and plant odour in the olfactory pathway of a male moth. PloS One, 7, e33159. (doi:10.1371/journal.pone.0033159)

Dumenil, C., Judd, G.J., Bosch, D., Baldessari, M., Gemeno, C. & Groot, A.T. (2014).Intraspecific variation in female sex pheromone of the codling moth Cydia pomonella. Insects, 5, 705–721. (doi:10.3390/insects5040705)

Ehrlich, P.R. & Raven, P.H. (1964).Butterflies and plants: a study in coevolution. Evolution, 18, 586–608. (doi:10.2307/2406212)

El-Sayed, A.M. (2023).The pherobase: Database of insect pheromones and semiochemicals. Available at: http://www.pherobase.com.Last accessed 23 August 2023.

Emelianov, I., Dres, M., Baltensweiler, W. & Mallet, J. (2001).Host-induced assortative mating in host races of the larch budmoth. Evolution, 55, 2002–2010. (doi:10.1111/j.0014-3820.2001.tb01317.x)

Fleischer, J., Pregitzer, P., Breer, H. & Krieger, J. (2018).Access to the odor world: olfactory receptors and their role for signal transduction in insects. Cell. Molec. Life Sc., 75, 485–508. (doi:10.1007/s00018-017-2627-5)

Gadenne, C., Barrozo, R.B. & Anton, S. (2016).Plasticity in insect olfaction: to smell or not to smell? Annu. Rev. Entomol., 61, 317–333. (doi:10.1146/annurev-ento-010715-023523)

Gonzalez, F., Witzgall, P., Walker, W.B. (2017).Antennal transcriptomes of three tortricid moths reveal putative conserved chemosensory receptors for social and habitat olfactory cues. Sci. Rep., 7, 41829 (doi:10.1038/srep41829)

Gonzalez, F., Borrero-Echeverry, F., Josvai, J.K., Strandh, M., Unelius, C.R., Toth, M. et al. (2020).Odorant receptor phylogeny confirms conserved channels for sex pheromone and host plant signals in tortricid moths. Ecol. Evol., 10, 7334–7348. (doi:10.1002/ece3.6458)

Ha, T. S. & Smith, D. P. (2022).Recent insights into insect olfactory receptors and odorant-binding proteins. Insects, 13, 926. (doi:10.3390/insects13100926)

Hansson, B.S. & Anton, S. (2000).Function and morphology of the antennal lobe: new developments. Annu. Rev. Entomol., 45, 203–231. (doi: 10.1146/annurev.ento.45.1.203)

Hatano, E., Saveer, A.M., Borrero-Echeverry, F., Strauch, M., Zakir, A., Bengtsson, M. et al. (2015).A herbivore-induced plant volatile interferes with host plant and mate location in moths through suppression of olfactory signalling pathways. BMC Biol., 13, 75 (doi:10.1186/s12915-015-0188-3) [50] [SLUPUb]

Janz, N. (2011).Ehrlich and Raven revisited: mechanisms underlying codiversification of plants and enemies. Annu. Rev. Ecol. Evol. Syst., 42, 71–89. (10.1146/annurev-ecolsys-102710-145024)

Knight, A.L. & Light, D.M. (2001).Attractants from Bartlett pear for codling moth, Cydia pomonella (L.), larvae. Naturwiss., 88, 339–42. (doi:10.1007/s001140100244)

Knight, A.L, Light, D.M., Judd, G.J.R. & Witzgall P. (2018).Pear ester - from discovery to delivery for improved codling moth management. In: Roles of Natural Products for Biorational Pesticides in Agriculture, eds. Beck, J.J., Rering, C.C. & Duke, S.O. (eds.). ACS Symposium Series 1294:83–113. (doi:10.1021/bk-2018-1294.ch008)

Kromann, S.H., Saveer, A.M., Binyameen, M., Bengtsson, M., Birgersson, G., Hansson, B.S. et al. (2015).Concurrent modulation of neuronal and behavioural olfactory responses to sex and host plant cues in a male moth. Proc. R. Soc. B, 282, 20141884. (doi:10.1098/rspb.2014.1884)

Landolt, P.J. & Phillips, T.W. (1997).Host plant influences on sex pheromone behavior of phytophagous insects. Annu. Rev. Entomol., 42, 371–391. (doi:10.1146/annurev.ento.42.1.371)

Light, D. M. & Knight, A. (2005).Specificity of codling moth (Lepidoptera: Tortricidae) for the host plant kairomone, ethyl (2E,4Z)-2,4-decadienoate: field bioassays with pome fruit volatiles, analogue, and isomeric compounds. J. Agric. Food Chem., 53, 4046–4053. (doi:10.1021/jf040431r)

Light, D.M., Knight, A.L., Henrick, C.A., Rajapaska, D., Lingren, B., Dickens, J.C. et al. (2001).A pear-derived kairomone with pheromonal potency that attracts male and female codling moth, Cydia pomonella (L.). Naturwiss., 88, 333–338. (doi:10.1007/s001140100243)

Ljunggren, J., Borrero-Echeverry, F., Chakraborty, A., Lindblom, T.U., Hedenström, E., Karlsson, M. et al. (2019).Yeast volatomes differentially effect larval feeding in an insect herbivore. Appl. Environm. Microbiol., 85, e01761–19 (doi:10.1128/AEM.01761-19)

Maan, M.E. & Seehausen, O. (2011).Ecology, sexual selection and speciation. Ecol. Lett. 14, 591–602. (doi:10.1111/j.1461-0248.2011.01606.x)

Makranczy, G., Tóth, M., Chambon, J.-P., Unelius, C.R., Bengtsson, M. & Witzgall, P. (1998).Sex pheromone of pear moth, Cydia pyrivora Danil. (Lepidoptera, Tortricidae). BioControl, 43, 339–344. (doi:10.1023/A:1009983613234)

Malausa, T., Bethenod, M.T., Bontemps, A., Bourguet, D., Cornuet, J.M. & Ponsard, S. (2005).Assortative mating in sympatric host races of the European corn borer. Science, 308, 258–260. (doi:10.1126/science.1107577)

Mallet, J., Meyer, A., Nosil, P., & Feder, J. L. (2009).Space, sympatry and speciation. J. Evol. Biol., 22, 2332–2341. (doi:10.1111/j.1420-9101.2009.01816.x)

McNair, C., Gries, G. & Gries, R. (2000).Cherry bark tortrix, Enarmonia formosana: Olfactory recognition of and behavioral deterrence by nonhost angio-and gymnosperm volatiles. Journal of Chemical Ecology, 26(4), 809–821. (doi: 10.1023/A:1005443822030)

Merrill, R.M., Wallbank, R.W.R., Bull, V., Salazar, P.C.A., Mallet, J., Stevens, M. & Jiggins, C.D. (2012).Disruptive ecological selection on a mating cue. Proc. R. Soc. B, 279, 4907–4913. (doi:10.1098/rspb.2012.1968)

Namiki, S., Iwabuchi, S. & Kanzaki, R. (2008).Representation of a mixture of pheromone and host plant odor by antennal lobe projection neurons of the silkmoth Bombyx mori. J. Comp. Physiol. A, 194, 501–515. (doi:10.1007/s00359-008-0325-3)

Pelozuelo, L., Malosse, C., Genestier, G., Guenego, H. & Frerot, B. (2004).Host-plant specialization in pheromone strains of the European corn borer Ostrinia nubilalis in France. J. Chem. Ecol., 30, 335–352. (doi:10.1023/B:JOEC.0000017981.03762.ed)

Pelozuelo, L., Meusnier, S., Audiot, P., Bourguet, D. & Ponsard, S. (2007).Assortative mating between European corn borer pheromone races: beyond assortative meeting. PLoS One, 2, e555. (doi:10.1371/journal.pone.0000555)

Phillips, P.A. & Barnes, M.M. (1975).Host race formation among sympatric apple, walnut, and plum populations of the codling moth, Laspeyresia pomonella. Ann. Entomol. Soc. Am. 68, 1053-1060. (doi:10.1093/aesa/68.6.1053)

Pregitzer, P., Schubert, M., Breer, H., Hansson, B. S., Sachse, S. & Krieger, J. (2013).Plant odorants interfere with detection of sex pheromone signals by male Heliothis virescens. Front. Cell. Neurosc., 6, 42. (doi:10.3389/fncel.2012.00042)

Priesner, E. & Baltensweiler W. (1987).Studien zum Pheromon-Polymorphismus von Zeiraphera diniana Gn. (Lep., Tortricidae). J. Appl. Ent., 104, 234–256. (doi:10.1111/j.1439-0418.1987.tb00545.x)

Ramdya, P. & Benton, R. (2010).Evolving olfactory systems on the fly. Tr. Genetics, 26, 307–316. (doi:10.1016/j.tig.2010.04.004)

Rebar, D. & Rodriguez, R.L. (2015).Insect mating signal and mate preference phenotypes covary among host plant genotypes. Evolution, 69, 602–610. (doi:10.1111/evo.12604)

Reddy, G.V. & Guerrero, A. (2004).Interactions of insect pheromones and plant semiochemicals. Tr. Plant Sc., 9, 253–261. (doi:10.1016/j.tplants.2004.03.009)

Ricci, B., Franck, P., Bouvier, J. C., Casado, D. & Lavigne, C. (2011).Effects of hedgerow characteristics on intra-orchard distribution of larval codling moth. Agric. Ecosyst. Environm., 140, 395–400. (doi:10.1016/j.agee.2011.01.001)

Ritchie, M.G. (2007).Sexual selection and speciation. Annu. Rev. Ecol. Evol. Syst. 38, 79–102. (doi:10.1146/annurev.ecolsys.38.091206.095733

Rosenthal, G.G. (2017).Mate choice: the evolution of sexual decision making from microbes to humans. Princeton University Press, New Jersey, pp. 648.

Safran, R.J., Scordato, E.S.C., Symes, L.B., Rodriguez, R.L. & Mendelson, T.C. (2013).Contributions of natural and sexual selection to the evolution of premating reproductive isolation: a research agenda. Tr. Ecol. Evol., 28, 643–650 (doi:10.1016/j.tree.2013.08.004)

Schluter, D. (2009).Evidence for ecological speciation and its alternative. Science, 323, 737–741. (doi:10.1126/science.1160006)

Scordato, E.S., Symes, L.B., Mendelson, T.C. & Safran, R.J. (2014).The role of ecology in speciation by sexual selection: a systematic empirical review. J. Hered., 105, 782–794. (doi:10.1093/jhered/esu037)

Servedio, M.R., Van Doorn, G.S., Kopp, M., Frame, A.M. & Nosil, P. (2011).Magic traits in speciation: ‘magic’ but not rare? Tr. Ecol. Evol., 26, 389–397. (doi:10.1016/j.tree.2011.04.005)

Smadja, C. & Butlin, R.K. (2009).On the scent of speciation: the chemosensory system and its role in premating isolation. Heredity, 102, 77–97. (10.1038/hdy.2008.55)

Trona, F., Anfora, G., Bengtsson, M., Witzgall, P., and Ignell, R. (2010).Coding and interaction of sex pheromone and plant volatile signals in the antennal lobe of the codling moth Cydia pomonella. J. Exp. Biol. 213, 4291–4303. (doi:10.1242/jeb.047365)

Trona, F., Anfora, G., Balkenius, A., Bengtsson, M., Tasin, M., Knight, A., Janz, N., Witzgall, P., and Ignell, R. (2013).Neural coding merges sex and habitat chemosensory signals in an insect herbivore. Proc. R. Soc. B 280, 20130267. (doi:10.1098/rspb.2013.0267)

Wan, F., Yin, C., Tang, R., Chen, M., Wu, Q., Huang, C. et al. (2019).A chromosome-level genome assembly of Cydia pomonella provides insights into chemical ecology and insecticide resistance. Nature Comm., 10, 4237. (doi:10.1038/s41467-019-12175-9)

Wang, F., Deng, J., Schal, C., Lou, Y., Zhou, G., Ye, B. et al. (2016).Non-host plant volatiles disrupt sex pheromone communication in a specialist herbivore. Sci. Rep., 6, 32666. (doi:10.1038/srep32666)

Walker, W.B., Gonzalez, F., Garczynski, S.F. & Witzgall, P. (2016).The chemosensory receptors of codling moth Cydia pomonella - expression in larvae and adults. Sci. Rep., 6, 23518. (doi:10.1038/srep23518)

Witzgall, P., Bengtsson, M., Unelius, C.R. & Löfqvist J. (1993).Attraction of pea moth Cydia nigricana F. (Lepidoptera: Tortricidae) to female sex pheromone (E,E)-8,10-dodecadien-1-yl acetate, is inhibited by geometric isomers (E,Z), (Z,E) and (Z,Z). J. Chem. Ecol., 19, 1917–1928. (doi:10.1007/BF00983796)

Witzgall, P., Chambon, J.-P., Bengtsson, M., Unelius, C.R., Appelgren, M., Makranczy, G. et al. (1996).Sex pheromones and attractants in the Eucosmini and Grapholitini (Lepidoptera, Tortricidae). Chemoecology 7:13–23. (doi:10.1007/BF01240633)

Witzgall, P., Bäckman, A.-C., Svensson, M., Koch, U.T., Rama, F., El-Sayed, A. et al. (1999).Behavioral observations of codling moth, Cydia pomonella, in orchards permeated with synthetic pheromone. BioControl, 44, 211–237. (doi:10.1023/A:1009976600272)

Witzgall, P., Ansebo, L., Yang, Z., Angeli, G., Sauphanor, B. & Bengtsson, M. (2005).Plant volatiles affect oviposition by codling moths. Chemoecology, 15, 77–83. (doi:10.1007/s00049-005-0295-7)

Witzgall, P., Stelinski, L., Gut, L. & Thomson, D. (2008).Codling moth management and chemical ecology. Annu. Rev. Entomol., 53, 503–522. (doi:10.1146/annurev.ento.53.103106.093323)

Witzgall, P., Trematerra, P., Liblikas, I., Bengtsson, M. & Unelius, C.R. (2010).Pheromone communication channels in tortricid moths: lower specificity of alcohol vs. acetate geometric isomer blends. Bull. Entomol. Res., 100, 225–230. (doi:10.1017/S0007485309990186)

Witzgall, P., Proffit, M., Rozpedowska, E., Becher, P.G., Andreadis, S., Coracini, M. et al. (2012).“This is not an apple” - yeast mutualism in codling moth. J. Chem. Ecol., 38, 949–957 (doi:10.1007/s10886-012-0158-y)

Yang, C., Cheng, J., Lin, J., Zheng, Y., Yu, X. & Sun, J. (2022).Sex pheromone receptors of lepidopteran insects. Front. Ecol. Evol., 10, 797287. (doi:10.3389/fevo.2022.797287)

Zakir, A., Sadek, M.M., Bengtsson, M., Hansson, B.S., Witzgall, P. & Anderson, P. (2013).Herbivore-induced plant volatiles provide associational resistance against an ovipositing herbivore. J. Ecol., 101, 410–417 (doi: 10.1111/1365-2745.12041)

Zhang, Q.H. & Schlyter, F. (2003).Redundancy, synergism, and active inhibitory range of non-host volatiles in reducing pheromone attraction in European spruce bark beetle Ips typographus. Oikos, 101, 299–310. (doi:10.1034/j.1600-0706.2003.111595.x)

Zhang, Q.H. & Schlyter, F. (2004).Olfactory recognition and behavioural avoidance of angiosperm nonhost volatiles by conifer-inhabiting bark beetles. Agric. Forest Entomol., 6, 1–20. (doi:10.1111/j.1461-9555.2004.00202.x)

